# Frequency bands EEG Biomarkers for Dementia using Graph Neural Networks

**DOI:** 10.1101/2025.08.18.670945

**Authors:** Mohamed Radwan, Pedro G. Lind, Anis Yazidi

## Abstract

We introduce a simple and interpretable model for classification of electroencephalography (EEG) signals. Our focus essentially is on using deep learning to study how connectivity patterns that are integrated to classify the EEG signals and highlight the important discriminative features used by the model in predictions. In this study, we utilize the connectivity features across different frequency bands in multi edge Graph Neural Networks (GNN) and showed that edge features are complimentary. We use a simple GNN model to predict Frontotemporal Dementia (FTD) in EEG. Our model is capable of achieving average accuracy of approximately 76% using Leave-One-Subject-Out-subject for FTD predictions which are better than the baselines and comparable to State of the arts models. In this article, we study the importance of the connectivity edges, nodes and frequency bands in the prediction of the model, focusing in explainable AI methods through saliency maps to interpret the model both locally and globally. The Saliency maps highlight the importance of Occipital and anterior temporal regions in the prediction of FTD. Furthermore, our results highlight the importance of Alpha and Theta bands in the prediction of FTD. Our observations align with previous research done using classical statistical methods. We argue that there are complimentary information in each each connectivity feature and frequency band brain networks. The impacts of each connectivity metrics on the prediction of the model are quantified to highlight the complimentary information in each connectivity measure.

## 1 Introduction and background

Frontotemporal Dementia (FTD) is a brain disorders caused by progressive degeneration of the frontal and temporal lobes of the brain. These regions are important for personality, behavior, language, and executive functions. FTD Often occurs earlier than Alzheimer’s disease and the patients retain memory until later stages. The progression of FTD and Alzheimer’s disease remains poorly understood. One promising area for understanding the mechanism of FTD is the identification of reliable biomarkers through studying functional connectivity between different brain regions. Brain Connectivity has the capacity to enable the early detection of pathophysiological processes affecting brain networks. These patterns of connectomics can be noticed before clinical symptoms emerge and before structural alterations are visible in the brain [7]. Very early alterations of functional connectivity network may be detectable in Alzheimer subjects before the accumulation of beta-Amyloid plaques and Tau proteins [20]. Studying the processes that are happening in the frequency bands can also be useful in understanding the pathology in more details. This study is driven by the motivation of understanding this pathology connectomics may eventually be helpful in detection of cognitive declines at the early course of the disease.

To study the brain connectomics, Electroencephalography (EEG) can be useful tool. EEG has the potential to be a diagnostic tool because it is relatively cheap and non invasive. EEG has the ability to capture brain dynamics and activity with high temporal resolution. Several methods have been applied using EEG data to extract discriminative features for distinguishing patients with neurodegenerative pathologies from healthy individuals using statistical methods. Recently, an increasing number of studies on diagnostic tools using deep learning techniques.

Deep Learning methods have been viewed as black box models where it is usually hard to understand how and why the model predicts specific outcome. Furthermore, predictive mistakes can happen due to this nature. This raises concerns regarding interpretability which arises mainly because of the complexity of deep learning models. Deep Learning models is used for automatic extraction of salient features from EEG signals to be used or predictions. Particularly, the model learns features representation from EEG signals and relate to the clinical characteristics of neurodegenerative pathologies such as AD, Dementia and MCI. Therefore, there is a growing demand for enhancing the interpretability of these deep learning models. In particular, there is a need to understand why the model extracts these features and how it is related to output. In this study, we aim to study those extracted features in details which may be useful in obtaining useful biomarkers for FTD diagnosis.

Graph Neural Networks (GNNs) are specific types of Deep Learning models that integrate both the EEG electrode features and coupling between these electrodes in the predictive model. GNNs have been used in different ways where the adjacency matrix is built using the spatial distance between those electrodes. This is based on the connection that close electrodes are more connected than far electrodes. However, closer spatial relationship may not guarantee a closer functional relationship. In this case, the functional connectivity would be useful to capture the complex coupling between the EEG electrodes. Additionally, the spatial connection between electrodes are binarized so that the two electrodes are either connected or not connected which is not optimal. The other way is using functional connectivity to build an adjacency matrix with continuous variables. Considering connectivity has the advantage of being better suited to study brain disorders [25], since it weights the level of connectivity instead of attributing binary labels. Typically the adjacency, in this case, is essentially a weighted fully connected matrix.

The study [5] used graph attention networks for generic classification of EEG without providing an interpretation. Another Graph based model model (DGCNN) was introduced by [25], that can learn dynamically the adjacency matrix that describes the connection between electrodes to be used for the task of emotion recognition. In [25], the model learns the relationships of the various nodes dynamically through the training of the model. In [31], the authors develop Graph Input layer attention convolutional network for classification of EEG data using Pearson correlation as adjacency matrix. The experiment is done on depression EEG dataset. The study [31] used extracted linear features from EEG (activity, mobility complexity and power spectral density) to be used as node features. They used the weight matrix in the input layer to find some edges and nodes that played an important role in depression recognition.

GNNs with explanation was developed by [30] that is based on Graph Convolutional Network using Chebyshev polynomial [4] to detect Parkinson diseases in EEG. The authors build an interesting framework to qualify the importance of nodes and edges in the decision of the model and draw conclusion about the abnormal connectivity. Those abnormal connectivity can be used as a biomarker in detection of Parkinson pathology. The study is complimented by the usage of microstates to understand the evolving dynamics of the brain. However this study lacks the usage of different frequency bands which could be useful to draw conclusion about the brain processes in the different bands. The study by [21] used Chebyshev polynomial based Graph Neural Network for abnormality classifications of EEG but this study lacks the usage of different frequency bands.

Interpretation methods have been used in other based on a Convolutional Neural network CNN to study mild cognitive impairments (MCI). In [27], the authors used CNN to classify two EEG MCI datasets and compare the performances of using four different frequency hands to the full frequency EEG images. They conclude and interpreting results that show that the Beta band (15 − 30 Hz) is the most important in the model decision as it gives the best performances in comparison with other frequency bands.

Another CNN based model is used by [16] to classify TUH [19] abnormal EEG datasets and compare the performances of using five different frequency hands to the full frequency EEG images. They conclude and interpreting results that show that the Delta band (0.5 − 4 Hz) is the most important in the model decision. Despite that the methodology is interesting and the study uses large dataset with rich content, the dataset are annotated as abnormal and normal classification. The study is limited in drawing a conclusion about certain pathological diseases. However, the study by [16] remains a general framework to be used for other specific datasets.

Rich Frequency content of the EEG signals can be further explored. In this study, we utilize the frequency content of the data in the predictive model and explainability. Understanding how these bands are related to advanced brain functions (e.g. learning, memory) can be helpful for both explainability and performances [9]. This also could be useful in understanding the FTD pathology and its mechanisms. The most relevant method to our study is the one proposed by [1]. The authors used Graph Neural Networks model for predictions of Alzheimer and Epilepsy cases with studying the feature importance used for Alzheimer predictions on Alzheimer/MCI dataset by [6]. The authors concluded the importance of Alpha waves in prediction of Alzheimer. In terms of the brain regions, occipital (O1) and prefrontal (Fp1) areas are the most important in prediction of Alzheimer. The study [1] focuses more on the model development. Here, we focus on the interpretation of the model and the extracted salient features used for the predictions. Furthermore, we use the method to study the Front temporal Dementia pathology.

Driven by the assumption that Deep Learning methods are capable of extracting the hidden patterns in the EEG data that are used for predictions, we aim to interpret the hidden extracted features by the Deep Learning model and understand it from neuroscientific perspective. We validate those observations with the findings in research in order to draw conclusion about the FTD pathology.

Our contribution in this study are:

– Providing a simple and competitive framework to be used for collectively interpreting the graph neural networks model in the classification of the EEG data. In the used case, we will use the model for the classification task of Frontotemporal Dementia vs Control (FTD vs CN).
– Explaining the predictions of the model using feature saliency maps to highlight the importance of the frequency bands, edges and brain regions in the classification of EEG. The heat maps of features are used to understand about the underlying FTD disorder that were recorded in the EEG which may serve as a biomarker for the disease.

The format of this study is as follows: first, we build a frequency based graph neural networks for binary classification the EEG data into FTD vs CN. Second, we use saliency maps to quantify the contributions of each features in the model output. We study the edge features and frequency bands using the model to draw conclusions about the complimentary information and the contributions of features in the predictions. Finally, we resort back to the data to study these contributing features and validate those findings with previous neuroscience research.

## 2 Data and Methods

### 2.1 Dataset

This dataset [15] has resting state-closed eyes EEG recordings from 88 subjects at the AHEPA University Hospital, Aristotle University of Thessaloniki in Greece. In this dataset, 36 of the subjects were diagnosed with Alzheimer’s disease (AD), 23 were diagnosed with Frontotemporal dementia (FTD), and 29 were healthy controls (CN). Here, we used the FTD and CN subjects. The EEG electrodes are 19 on the scalp and 2 reference electrodes. The sampling rate was 500 Hz. In this study, we used the preprocessed dataset by the authors [15]. According to [15], The signals were re-referenced to the A1-A2 after applying a Butterworth band-pass filter with a frequency range of 0.5 to 45 Hz. This is followed by ASR routine and ICA algorithms to remove large amplitude and eye artifacts. Informed consent was obtained by [15] from all subjects by the AHEPA University Hospital Aristotle University of Thessaloniki in Greece.

### 2.2 Used Architecture

Using multichannels EEG dataset, we extract node features for the graph *G* = (𝒱,*ℰ*). Here, *V* is the number of vertices and *E* is the set of edges. Each node of the graph *V* has an input features. The features are calculated for each frequency band. Examples of node features are entropy, mobility, complexity. Calculating the adjacency matrices is explained in details in section 2.5. In our methods, we formulated the problem as graph classification. We adopt the Transformer Convolution operator which generates node embedding to be used for graph classification via global mean pooling.

Transformer Convolution operator employs attention in message passing is based on weighting which node embeddings are important through GNNs message passing. This can be one single attention weight for each pair of nodes. Transformer Convolution operator [23] employs multi-head attention instead of one single attention for each pair of nodes before aggregating the embeddings. The aggregation of attention graph networks with a single attention head is as the following

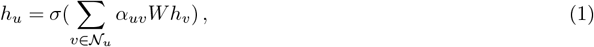

where *σ* is sigmoid activation, *h*_*v*_ and *h*_*u*_ are neighbouring (*v*) and target (*u*) nodes embeddings, *W* is the projection matrix of convolutional operator and *α*_*uv*_ are normalized attention scores. To extend this with multihead attention for *K* neighbors as said in Transformer Convolution operator, we have:

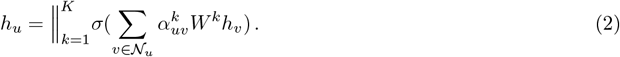

The output from this operator are node embeddings in a graph. So we use global mean pooling that returns batch-wise graph level embedding *h*_*g*_ by mean aggregation of the node features across the node dimension for *n* is the number of nodes, namely

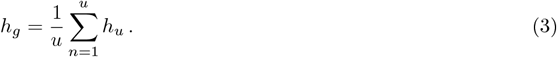

The network architecture is shown in Fig 1. This network is trained using multi edges graph. In this study, we extract the frequency bands from the EEG data. The module explained here is the main building block for the predictive model used in this study. Decomposing EEG signals will be explained in the next section 2.3.

**Fig. 1:**
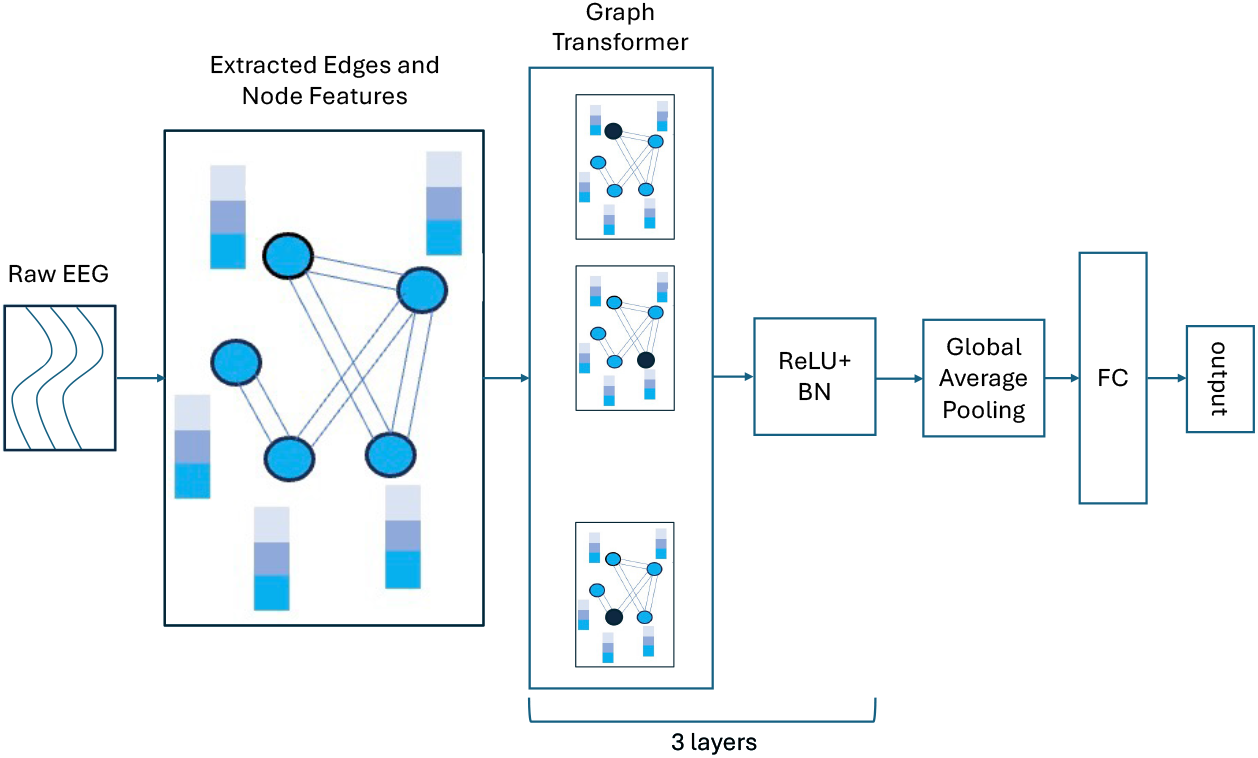
Used Model Architecture. The abbreviation ReLU means rectified linear unit, BN means Batch Normalization and FC means Fully Connected layer. Raw EEG data is used to calculated edge and node features which generates a graph for each data sample. The graph is fed to the Graph Transformer operator followed by ReLU and BN layers. The Graph Transformer operator layer is repeated three times. The output of Graph Transformer operator is a node embedding where a global mean pooling layer is applied to generate a graph embedding before feeding to a Fully Connected layer (FC). The number of heads for the Graph Transformer operator is five

### 2.3 Frequency bands

The time domain signals are decomposed into frequency components before calculation of connectivity. The functional connectivity measures are calculated on the different frequency bands of EEG. The original data is converted using Time-frequency decomposition method using morlet wavelets. The five used frequency bands and their corresponding frequency ranges are: Delta (1-4 Hz), Theta (4-8 Hz), Alpha (8-13 Hz), Beta (13-30 Hz) and Gamma (30-45 Hz).

Furthermore, We calculate the nodes features and edges for the different frequency bands before feeding them to the model. For each frequency band, there is a graph node and edge features. The calculation of node and edge features is explained in the sections 2.4 and 2.5.

### 2.4 EEG Extracted Features

The window size in the experiments here is 30 seconds which is the same window size used by [14] on this dataset for fair comparison. The used model is trained on these windows and the prediction task is window or epoch based classifications. The extracted saliency maps are computed from those windows. With the aim to train a classifier to recognize time-varying dynamics, a sliding window is used allowing the increment of samples to help the model in capturing dynamics that evolves over time. The overlap between windows is half the window size which is 15 seconds. According to [8], the selection of window size is not straightforward. If a window is too short, the stimulus that induced responses may not have been fully unfolded during the window. On the other hand, long windows might fail to capture salient features or mix different states.

Hand crafted features are extracted from the EEG data. The extracted features are differential entropy, Spectral entropy, Permutation entropy, entropy power, approximate entropy, Hjorth mobility and complexity, approximate entropy, fractal density and power spectrum density. Those features are complimented with mean, standard deviation and kurtosis of the signal. Those features are extracted on each of the frequency bands.

### 2.5 Functional Connectivity

Three different functional Connectivity metrics are used in this study to quantify the coupling relationship between the EEG electrodes. The connectivity metrics are Spectral Coherence, Phase Locking Value and Phase Locking Index.

#### Spectral Coherence

Coherence is used to measure the magnitude squared coherence estimate, *COH* using Welch’s method as:

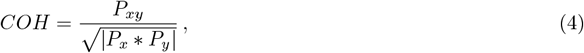

where *P*_*x*_ and *P*_*y*_ are power spectral density estimates of *X* and *Y*, and *P*_*xy*_ is the cross spectral density estimate of *X* and *Y*.

#### Phase Locking Value (PLV)

Phase Locking Value (PLV) or Phase synchronization [10] can be used to measure functional connectivity between two signals based on the phase. This means that PLV considers consistent phase differencing between the two signals while ignoring the amplitudes. This is can be useful in case the amplitudes magnitudes of the two signals are not comparable which is the aim in this study.

The PLV is given as the average phase difference between two signals as:

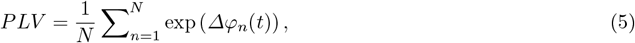

where *Δφ*_*n*_(*t*) is the phase difference between the two signals at time *t* and *N* is the number of time points. The Phase difference is calculated using Hilbert transform which computes the instantaneous phase of the signal which gives value between is a value between 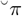 and *π*. PLV takes values between 0 and 1 with 0 reflecting that there is no phase synchrony and 1 where the relative phase is identical and consistent.

#### Phase Locking Index (PLI)

Similar to PLV, Phase Locking Index (PLI) [26] quantifies the phase synchronization of two signals. However, PLV can have large inflated value as a result of a single source contributing to both signals due to volume conduction in the channels. PLI is used to tackle the the common source problem. PLI disregards the phase locking that is centered around 0 or *π* phase difference. It quantifies the asymmetry of the distribution of relative phase around zero using the sign function of phase differences between the two signals and is defiend as:

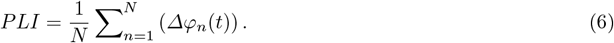

PLI takes values between 0 and 1. The value is 0 if the distribution of relative phase is symmetric about 0or *π*. Three types of functional connectivities are used for each frequency band. This results in 15 edge features. The adjacency matrix is of the 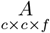 where *f* are the edge features and *c* is the EEG channel.

### 2.6 Explainable AI (XAI) Methods

Saliency maps are developed by initializing a weight tensor that is differentiable. The used trained model will then assign higher weights to the most important nodes or edges that impacts the outcomes of the model. Saliency maps returns the gradient of the output with respect to the input. This approach can be understood as taking a first-order Taylor expansion of the network at the input, and the gradients are simply the coefficients of each feature in the linear representation of the model. The absolute value of these coefficients can be taken to represent feature importance [24]. Those values can be averaged over the whole dataset to generate global feature importance.

### 2.7 Experiments

In this experiment, the data are split using Leave One Out subject validation to compare to the baselines by [15]. LOSO validation is a robust and unbiased estimate of model performance especially for small dataset in the used case. In this experiment, we leave one EEG subject as a validation subject to be used for calculation of the performances and the rest of subjects are used for training. For each cross validation, the model is trained on the train data and tested on the LOSO subject. The accuracy for each LOSO subject is averaged over all the cross validation runs. The predictive task is binary classification of front temporal dementia vs Control (FTD vs CN).

In this experiments, we measure the metrics: Accuracy (Acc), Sensitivity (Sens), Specificity (Spec) and F1-score for evaluation. The evaluation metrics are calculated given the true positive (TP), true negative (TN), false positive (FP) and true negative (FN) as the following:

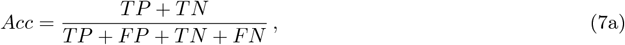

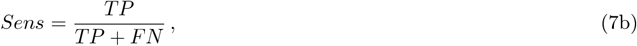

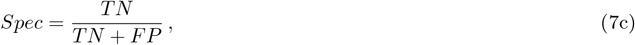

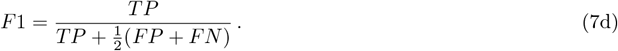

For choice of model hyperparameters, we used a small learning rate 0.00001 in order to avoid instability and divergence of the model. The models are trained for 35 number of epochs. After 35 epochs, the model starts to overfit. The used model architecture consists of three layers of graph convolutions. In this study, the prediction in this study is epoch or window based classification. This means that each window is binary classified to either CN or FTD. Since we use LOSO validation and knowing that each LOSO subject belongs to either of the classes. Therefore, we take majority voting of the predictions and generalize it over the whole LOSO subject epochs. This means that if the majority of epochs in a LOSO subject are classified as FTD, the whole subject epochs are classified as FTD. The number of parameters in this model architecture is 281, 954.

## 3 Results

The used model achieved weighted average LOSO accuracy of 76.06% in the prediction of FTD as reported in Table 1. The reported accuracy is better than in the baselines in [15]. The accuracy of the proposed model is nearly 4% better than the best performing baselines.

**Table 1:** Comparison between our model and several baselines from [15] for the task of classification of front temporal dementia (FTD) and healthy control (CN). The shown results weighted average of LOSO cross validation performances.

| FTD vs CN | Accuracy | Sensitivity | Specificity | F1 Score |
| --- | --- | --- | --- | --- |
| kNN | 67.34% | 59.67% | 76.13% | 70.81% |
| SVM | 70.14% | 62.41% | 75.98% | 68.32% |
| MLP | 73.12% | 63.00% | 78.63% | **72.82%** |
| LightGBM | 72.43% | 61.13% | **80.74%** | 66.31% |
| Random Forest | 72.01% | 72.32% | 72.32% | 67.32% |
| Our Model | **76.06%** | **78.95%** | 74.89% | 65.44% |

The used model shows an improvement in sensitivity. The improvements are nearly 6% than the best performing baseline. This is valuable, given the sensitivity of the case study, as it means that the model is better at increasing the true positive subjects with FTD. This means that the risk of missed True Positive patients is reduced. However, The higher sensitivity comes with lower specificity as shown in the Table 1.

For further comparison, we compare our reported results (Table 2) with other models using Deep Learning architectures. DICE-net [14], which is the state of the art performance on this data, achieves the best performances except in the sensitivity which our model is capable of achieving the best performance. However, the accuracy of our model is closely comparable to best performing model with less than 1% difference. It is also noticed that other State Of The Art EEG models are showing significant lower performances in both the problems. It is noted that our model is achieving better performances compared to DeepConvNet, EEGNet [11] and EEGNETSSVEP in the used case.

**Table 2:** Comparison between our model and several Deep Learning models from [14] for the task of classification of front temporal dementia (FTD) and healthy control (CN). The shown results weighted average of the performances of 10 cross validations.

| FTD vs CN | Accuracy | Sensitivity | Specificity | F1 Score |
| --- | --- | --- | --- | --- |
| DeepConvNet | 64.21% | 62.41% | 37.05% | 60.20% |
| EEGNet [11] | 46.00% | 42.20% | 57.46% | 43.65% |
| EEGNetSSVEP [29] | 61.46% | 53.51% | 75.00% | 52.43% |
| DICE-net | **74.96%** | 60.62% | **78.63%** | **62.27%** |
| Our Model | 74.16% | **89.09%** | 70.69% | 56.49% |

In the next section, we discuss the salient features that are used by the models for predictions. The following sections visualize and highlight the important features used by the model in the predictions. More specifically, we show the saliency maps for important nodes, edges and frequency bands.

### 3.1 Node Saliency Maps

Fig 2 shows the importance of the nodes in the used classification problem. The heatmaps show the relative importance of each node in the brain network with respect to the minimum and maximum of the node importance. By looking at the brain regions, the area of the electrodes *O*2, *T* 6, *T* 4, *T* 3 *P* 4, in the Alpha (8-13 Hz), Theta (4-8 Hz), Gamma (30-45 Hz) bands, are the most important nodes.

**Fig. 2:**
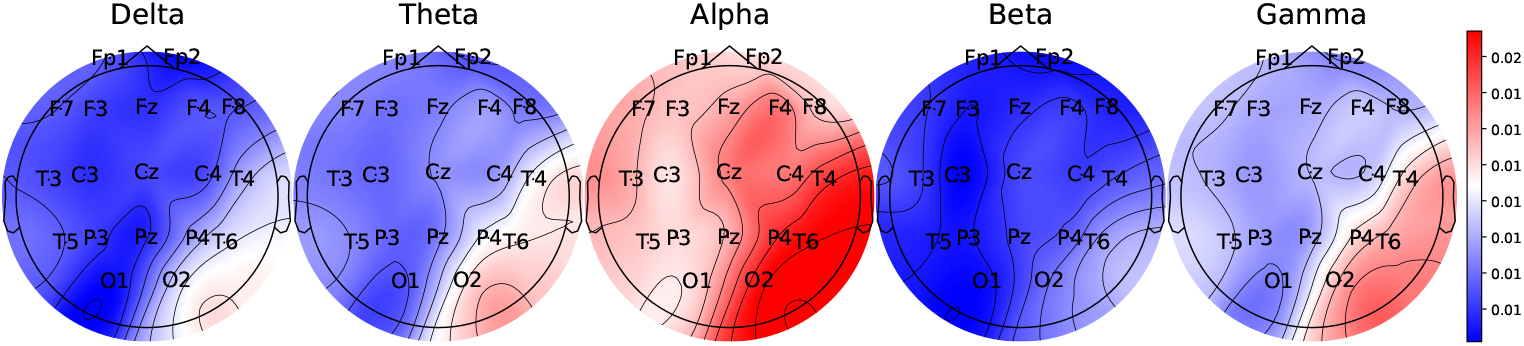
Importance of Nodes in the FTD vs CN classification using all node features of the LOSO subjects and the average of the importance of nodes.

### 3.2 Edges Saliency Maps

Fig 3 shows the importance of the edges in the prediction of FTD. Similar to the importance of nodes, the heatmaps show the relative importance of each edge with respect to the minimum and maximum values of in all the edges. It is noted that Theta band (4-8) is the most important band followed by Alpha and Gamma bands. PLV network has the highest impact on the model followed by Coherence. In the next section 3.3, we will go in depth about those important networks.

**Fig. 3:**
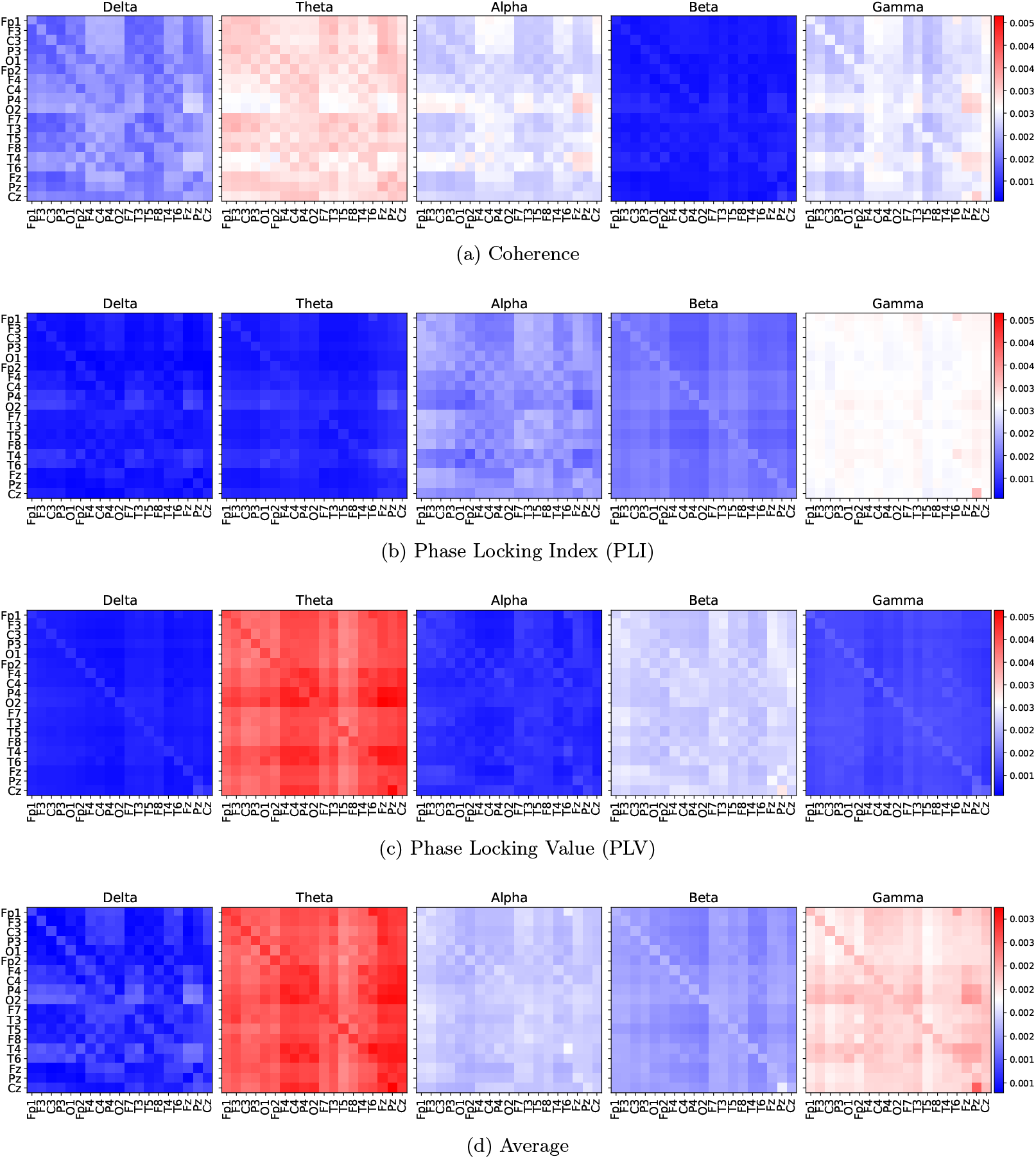
FTD vs CN classification relative Importance of edges for the different connectivity metrics for the different frequency bands.

### 3.3 Frontotemporal Dementia Brain Network Maps

From the previous section 3.2, it is noted that PLV in Theta band, Coherence in Theta, Alpha, Delta bands and PLI in Gamma band are highlighted as important networks that impact the model outcomes. Here, we look closely on each of the networks separately. The subfigure 4a shows the PLV in the theta band. It highlights semi-symmetric connections between the electrodes. The connection from and to the electrodes *O*2, *C*4, *P* 4, *P* 4 are highlighted with higher weight that is consistent with the observation from nodes importance in Fig 2 which shows that *O*2, *T* 6 and *P* 4 are most important nodes in the predictions of the model. In the subfigures 4c, 4d, 4f the connections the area which has the electrodes *O*2, *P* 4, *T* 4, *T* 6 and other electrodes *Fz, Pz, Fp*1, *F* 3, 01, *Fp*2 which belong to different brain regions. The connection between *Cz* and all other electrodes are highlighted the subfigures 4a, 4b, 4c, 4d, 4f. The connection between *Pz* and *Cz* is important in subfigures 4e and 4f.

**Fig. 4:**
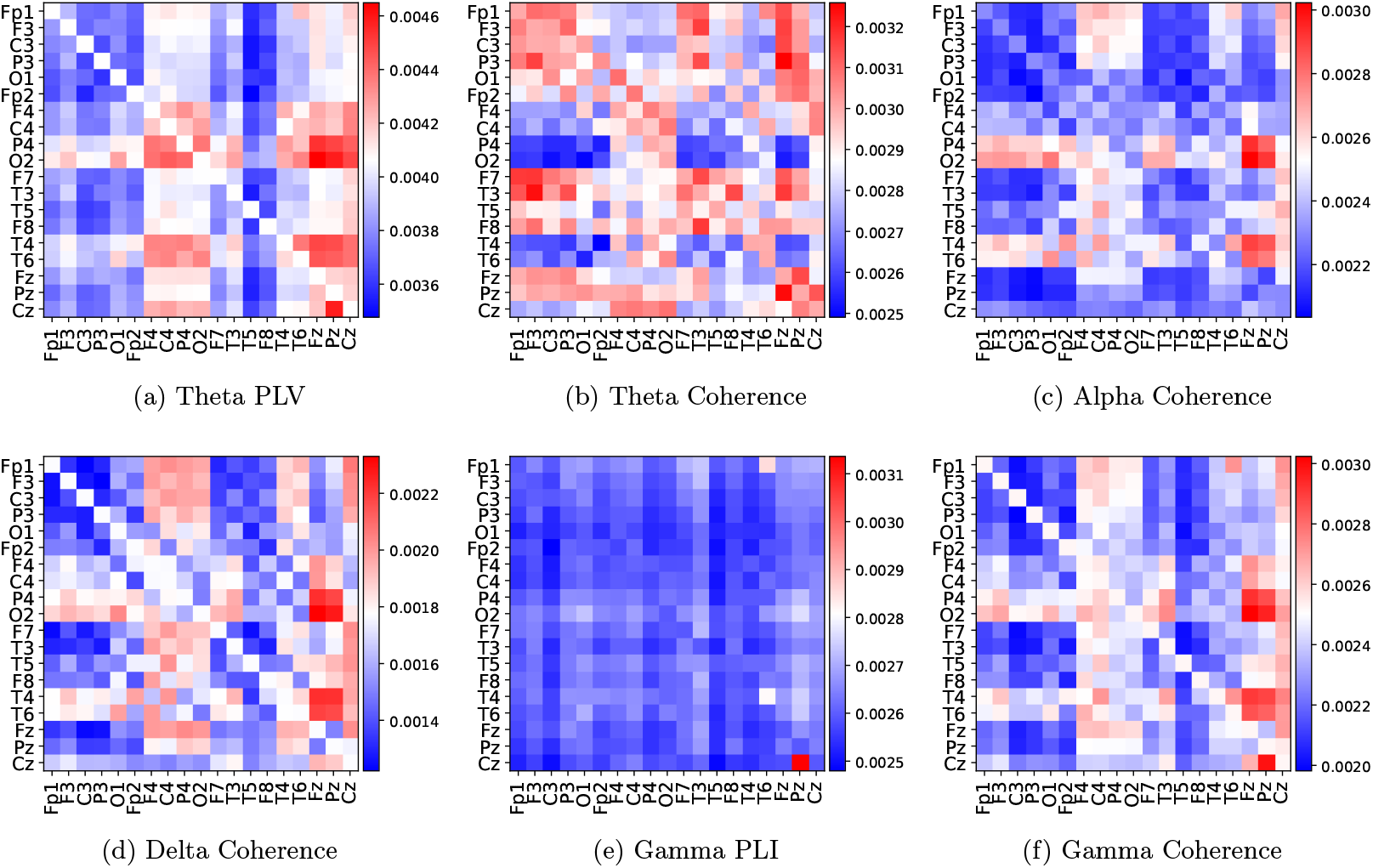
FTD vs CN classification relative Importance of edges for the most important network features of the different connectivity metrics for the different frequency bands. The plots are normalized to minimum and maximum of each plot to highlight most important edges.

## 4 Discussion

The proposed classifier is capable to achieve better performances than the baselines from [15] as shown in Table 1 and better than or comparable with state of the art models as shown in Table 2. The accuracy improvements are around 4% compared to the best baseline. The overall performance is higher is attributed to the higher sensitivity. High sensitivity means that more true positive findings for patients with a disease are predicted correctly which is critical in medical applications. This means that the proposed model have higher ability to to detect FTD disease. The higher sensitivity is also supported when comparing the proposed classifier with State Of The Art models such as DICE-net [14]. As sensitivity increases, specificity decreases which is a weakness of this model. Our proposed method shows lower specificity than the best baselines on used case.

Regarding The size of the proposed models, they have approximately 282 thousands parameters. These number of parameters are significantly lower than the number of parameters used in DICE-net which is 170.5 millions, according to [14]. This means that our models are more simple and efficient in general while maintaining competitive performances.

It is noted that the other deep learning models have lower performances than out model. That is because these models such as EEGNet and DeepConvNet rely on the raw amplitudes of the EEG signals which maybe noisy. The significant lower performances are reported in Table 2. Our proposed method has been more successful as it takes extracted hand crafted node features instead of learnable features through deep learning. Similarly, DICE-net [14] is successful as it takes Power spectral coherence and relative band power through morlet wavelet transform to be fed to the model. We argue that the amplitudes of the EEG signals maybe are not comparable due to presence of noise. In general, it is challenging to have the same experimental setting when placing EEG electrodes on the scalp and data recording. This makes signals not comparable in terms of amplitudes and very prone to noise which would make the scaling by removing the mean and dividing by the unit variance of the data not useful in the training process. Deep learning models will not be able to learn the relevant features as there is no clear patterns in the comparison between the raw amplitudes. We argue that to avoid this behaviour, we would need more data in which amount of data can compensate for the noise or using extracted features as we conduct our experiments in this study.

As stated earlier, our objective is the explanation of the extracted features used by the model for prediction of FTD. Our results highlight the importance of Alpha waves as discriminative features for the predictions. This observation is confirmed in saliency maps. Second most important bands are Gamma, Theta or Beta. Using the node features, it is noted that Alpha followed by Gamma, Theta and Beta bands features are the most impacteful features on the decision of the FTD vs CN classifier. This observation align with the study of [1] which conclude the importance of Alpha band in Alzheimer prediction. On the other hand, on the saliency of edge features, Theta bands are most important followed by Alpha or Gamma.

Several research studied the frequency bands importance in FTD disease with consistent evidences that abnormalities in EEG delta, theta, and alpha rhythms in FTD and Alzheimer. FTD generally exhibits decreased alpha power notably in frontal and temporal regions [13]. The Power Spectrum Density in was significantly reduced in alpha and beta bands for FTD subjects. The study [18] reports lower activity were observed in the alpha band for FTD group than Control in the orbital frontal and temporal lobe. FTD patients show increased theta power compared to controls [13], [17], [28].

### 4.1 Brain Regions

Occipital (*O*2 electrode) and left and right anterior temporal lobes (*T* 3 and *T* 4 electrodes) regions are highlighted as the most impactful region in the decision of both classifier. Occipital region significance is strongly evident in Alpha Band but also is highlighted in almost all used frequency bands. This results are consistent with [14] which is reported on the same data.

The extracted saliency maps separately for Node features and edge features. The saliency maps of the brain networks or the connectivity features confirm the observations from node features about the attention of the model to the Occipital and temporal lobes. This will be discussed in further details in the section 4.2.

Brain regions alterations happen in various neurodegenerative diseases. The study [12] highlights that healthy group showed a peak in activity at the frontal and Occipital regions that was reduced or absent from the mild Alzheimer group.

### 4.2 Brain Networks

Several interesting observation have been reported in the brain networks. The connections *O*2 and other electrodes are shown as important in the figures of connectivity saliency. The connections to and from the electrode *T* 3 and *T* 4 which are left and right anterior temporal lobes are also highlighted. The important connections between electrodes align with the important nodes that are mentioned earlier. Functional connectivity between the medial temporal lobe and the anterior-temporal system is reduced in the earliest stages of Alzheimer’s disease [2]. The studies [22] and [3] indicate that Medial temporal lobe connectivity networks show changes with higher age and in subjects with MCI and are related to cognition as well as future cognitive decline.

## 5 Conclusion and Future Directions

We introduce a model that achieves better than the baselines and comparable performances to the state of the art with significant lower number of trained parameters. In this study, we used the frequency bands connectivity and node features as input for classification of the data. First observation is the significance of Theta and Alpha waves in the prediction FTD in EEG data which aligns with prior research. The used saliency maps show consistent results for important nodes, bands and connectivity. Saliency maps and indicate that Occipital and temporal lobes specifically in the alpha band are the most important discriminative features from the EEG. Connectivity in the Occipital between *O*2 electrode in the Occipital region and other electrodes are highlighted as important.

The used Coherence, PLI and PLV metrics generates a symmetric connectivity between electrodes which still limited in terms of capturing the coupling between electrodes. One possible direction is to use directional connectivity such as Granger Causality which may capture more information and lead to better interpretation of the functional connectivity between electrodes. Additionally, A comparative analysis using different functional connectivity could be useful to further understanding of the connectivity for this pathology.

The used model takes hand crafted features as input which can be limiting in the case study. However, the use of raw time series features does not lead to convergence. This maybe is due to amount of noise in the data and limited size of the data. We argue that a data of a bigger size may lead to better performance when working with raw EEG data.

There are several limitations in this study. The first limitation is due to the challenge of the small data which makes the feature learning from raw EEG using deep learning challenging. It would be useful to extend the experiments to a large datasets which would lead to better generalization over broad subjects. Large and well curated data may compensate for the noise in the data. This is assuming that the noise is random and it does not exist with the same magnitude in all EEG electrodes in all data samples. The deep learning models can be more immune to the noise in a bigger data. This is a major limitations of the ability of Deep Learning methods given the limited size of available datasets in this domain and the complexity of the data collections and sharing.

The prediction task are using windows for classification. As each window is considered as a data sample. This does not take into account how those windows are connected dynamically to each other in the same subject which is suboptimal. A new model should be used that take in to account the dynamical relationship between windows in the same subjects. Dynamic graphs provide an possible direction to address this weakness.

## Acknowledgments

This work is part of the AI-Mind project. AI-Mind has received funding from the European Union’s Horizon 2020 research and innovation program (https://www.ai-mind.eu).

